# *Pseudomonas aeruginosa* cytochrome P450 CYP168A1 is a fatty acid hydroxylase that metabolizes arachidonic acid to the vasodilator 19-HETE

**DOI:** 10.1101/2021.10.19.465045

**Authors:** Brian C. Tooker, Sylvie E. Kandel, Hannah M. Work, Jed N. Lampe

## Abstract

*Pseudomonas aeruginosa* is a gram-negative opportunistic human pathogen that is highly prevalent in individuals with cystic fibrosis (CF). A major problem in treating CF patients infected with *P. aeruginosa* is the development of antibiotic resistance. Therefore, the identification of novel *P. aeruginosa* antibiotic drug targets is of the upmost urgency. The genome of *P. aeruginosa* contains four putative cytochrome P450 enzymes (CYPs) of unknown function that have never before been characterized. Analogous to some of the CYPs from *M. tuberculosis*, the *P. aeruginosa* CYPs may be important for growth and colonization of the CF patient’s lung. In this study, we cloned, expressed, and characterized CYP168A1 from *P. aeruginosa* and identified it as a sub-terminal fatty acid hydroxylase. Spectral binding data and computational modeling of substrates and inhibitors suggest that CYP168A1 has a large, expansive active site preferring long chain fatty acids and large hydrophobic inhibitors. Furthermore, metabolism experiments confirm that the enzyme is capable of hydroxylating arachidonic acid, an important inflammatory signaling molecule present in abundance in the CF lung, to 19-hydroxyeicosatetraenoic acid (19-HETE; *K_m_* = 41.1 µM, *V_max_* = 222 pmol/min/nmol P450), a potent vasoconstrictor which may play a role in the pathogen’s ability to colonize the mammalian lung. Metabolism of arachidonic acid is subject to substrate inhibition and is also inhibited by the presence of ketoconazole. This study points to the discovery of a new potential drug target that may be of utility in treating drug resistant *P. aeruginosa*.

## INTRODUCTION

*Pseudomonas aeruginosa* is a gram-negative opportunistic pathogen that is highly prevalent in individuals with cystic fibrosis (CF), a debilitating inherited disease (1–3). In CF patients with reoccurring *P. aeruginosa* infections, the organism is known to form intractable biofilms that lead to antibiotic resistance (4–7). Antibiotic failure can result in pneumonia which can be life-threatening in the respiratory compromised CF patient (2, 8). Moreover, chronic airway infection by *P. aeruginosa* significantly promotes lung tissue destruction, further compromising pulmonary function in individuals with CF. Given this, the need for characterization of new antibiotic targets and drugs that have the potential to inhibit biofilm formation in *P. aeruginosa* is urgently needed. The genome of *P. aeruginosa* strain PAO1 (UW) was first completely sequenced in 1999 (9) and serves as a reference genome for the organism (http://www.pseudomonas.com/). The genome of *P. aeruginosa* contains four putative cytochrome P450 (CYP) monooxygenase genes; designated CYP107S1, CYP168A1, CYP169A1, and CYP239A1(10). In bacteria, CYP enzymes are known to perform diverse functions, including: antibiotic synthesis (11–14), carbon source metabolism(15–17), detoxification(18, 19), and secondary metabolite production(20–22). Cytochrome P450 enzymes from the closely related obligate intracellular pathogen *M. tuberculosis* have been characterized and determined to be integral to the metabolism of fatty acids(23–26), cholesterol(27, 28) and even certain antibiotics(29, 30), all of which promote bacterial survival and growth under a variety of conditions. While the exact functions of the CYP enzymes identified in *P. aeruginosa* remain unknown, they have recently been implicated in both detoxification(18) and the oxidation of environmental contaminants (31). Further evidence links them to the metabolism of medium to long chain alkanes(32), suggesting their role as fatty acid and/or alkane hydroxylases. Indeed, a number of bacterial CYP enzymes are known to function as fatty acid hydroxylases and this role for CYPs seems to be common among the eubacteria(10), although this particular function has yet to be demonstrated for the *P. aeruginosa* CYP enzymes.

Many types of fatty acids are found within the environment of the lung and could possibly serve as carbon sources for *P. aeruginosa* as it attempts to establish a colony(8). However, certain fatty acids, including arachidonic acid and its metabolites, control inflammation and immune cell recruitment to the site of infection(33–35). Moreover, concentrations of arachidonic acid are significantly increased in the CF patient lung due to a metabolic defect(36, 37). An intriguing possibility is that the organism may be able to modify the immune response by oxidation of arachidonic acid and its metabolites, thereby making its local environment more hospitable for the parasite to grow and proliferate. Indeed, a secreted lipoxygenase from *P. aeruginosa*, designated LoxA, is capable of metabolizing arachidonic acid to 15-hydroxyeicosatetraenoic acid (15-HETE)(38), which has been demonstrated to down regulate the immune response *in vivo*(39), illustrating the capacity of *P. aeruginosa* to reduce the host’s ability to respond to infection through the metabolism of inflammatory mediators. More recent work elucidated the role of a soluble epoxide hydrolase to specifically reduce levels of the host pro-resolving lipid mediator, 15-epi lipoxin A4 (15-epi LXA4), thereby contributing to the prolongation of pulmonary inflammation and associated loss of lung function in patients with CF(40, 41). These studies raise the prospect that the modulation of anti-inflammatory lipid mediators may be a general strategy by which *P. aeruginosa* regulates the host–pathogen relationship to promote the colony establishment. Despite the evidence for the involvement of these bacterial enzymes in the regulation of inflammatory lipid mediators, the CYP enzymes of *P. aeruginosa* have never before been examined for their ability to metabolize physiologically important fatty acids, such as arachidonic acid and its derivatives.

Here, we report the characterization of the first CYP enzyme cloned from *P. aeruginosa*, CYP168A1(42). Our findings indicate that CYP168A1 is a fatty acid hydroxylase with a high affinity for long chain fatty acids, such as oleic and arachidonic acid. Additionally, CYP168A1 is capable of binding large azole inhibitors, including ketoconazole and miconazole. These results, and in silico modeling of ligands docked to the enzyme, suggest an enzyme with a large, expansive active site. Furthermore, we have characterized the hydroxylation pattern of arachidonic acid and lauric acid, a model fatty acid, establishing CYP168A1 as a medium to long chain fatty acid hydroxylase, attacking the carbon chain at the ω-1 and ω-2 positions. Moreover, it is capable of metabolizing arachidonic acid to 19-hydroxyeicosatetraenoic acid (19-HETE), a potent vasoconstrictor which may play an important role in the pathogen’s ability to colonize the mammalian lung. These results lay the groundwork for understanding the function of the CYP168A1 enzyme in this important human pathogen and its potential as a drug target.

## RESULTS

### Expression and Characterization of CYP168A1

CYP168A1 is the first CYP enzyme from *P. aeruginosa* PAO1 to be cloned, expressed, purified, and characterized in a soluble recombinant form(42). The protein was isolated and purified to homogeneity (>90%), resulting in a single band present on the denatured SDS-PAGE gel with a relative molecular weight of ∼48.5 kDa (Figure 1A). The purified protein exhibited a characteristic spectral absorption pattern for a heme containing protein, with a λ_max_ of 417 nm for the oxidized protein and 420 nm for the sodium dithionite reduced species (Figure 1B). Reduction of the CYP168A1 heme iron by sodium dithionite followed by carbon monoxide (CO) binding shifted the Soret peak to ∼450 nm, the signature of a cytochrome P450 enzyme (Figure 1C). The Soret peak at 450 nm grew with time, ultimately reaching a plateau and stabilizing after 20 min. Our initial expression experiment from 6 L of *E. coli* culture yielded a total of 315 nmoles of purified and soluble recombinant CYP168A1.

**Figure 1.**
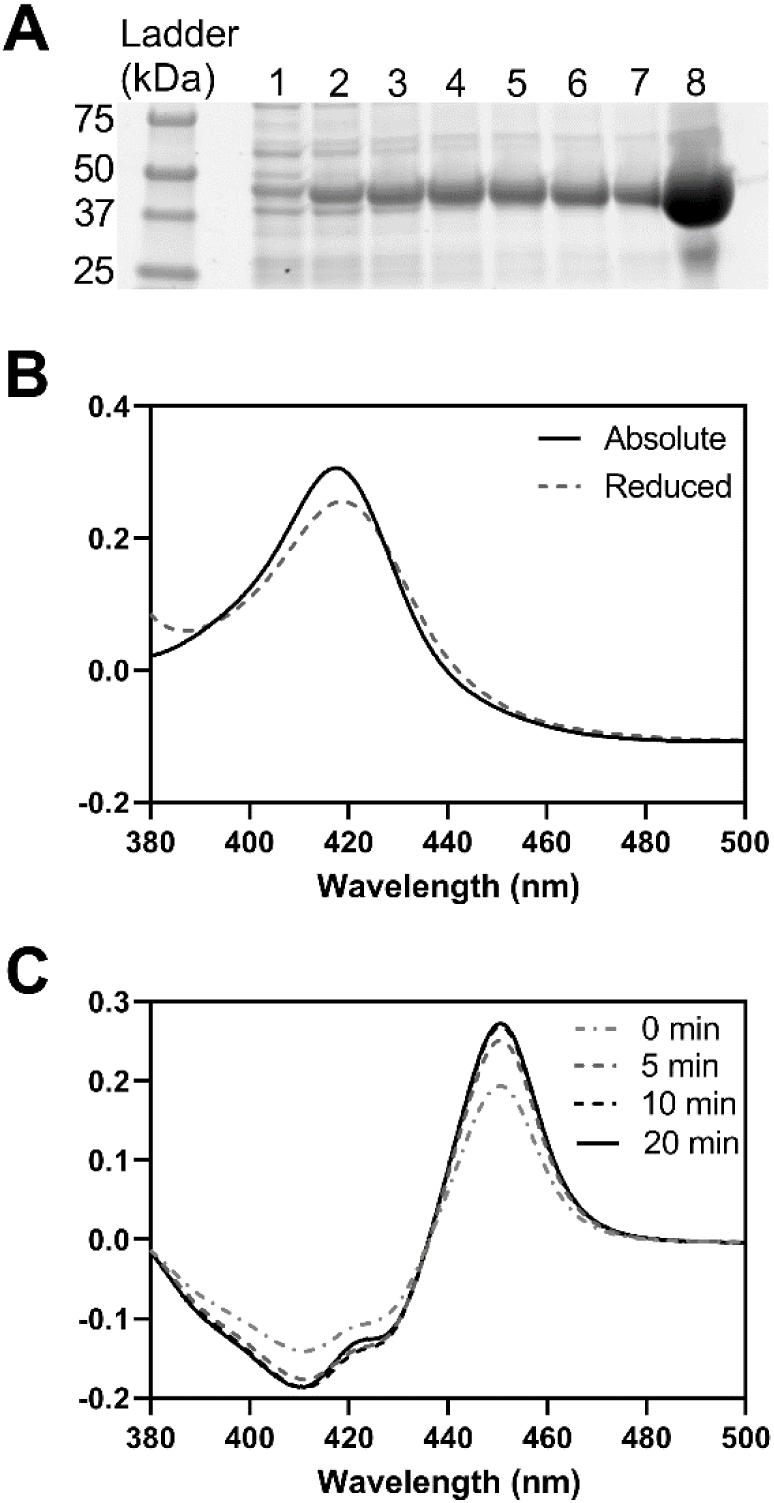
Expression and Spectral Absorption Characteristics of CYP168A1. (A) SDS-PAGE for the Ni-NTA fractions (# 1 to 8) of the purified recombinant CYP168A1 protein expressed with a C-terminal 4xHis-tag. (B) Oxidized and reduced absorption spectra of CYP168A1. (C) CO-binding difference spectra of the reduced CYP168A1 over a 20 min period.

### Ligand Binding to CYP168A1

In order to determine the chemical space occupied by ligands of CYP168A1, a series of spectral titrations were conducted with a variety of putative substrates and inhibitors, with selection criteria being based upon structural similarity to known cytochrome P450 ligands. Initial experiments focused on a variety of azole compounds, which are well known CYP inhibitors (Figure 2). All azole ligands examined elicited a typical Type II red shift of the Soret band to ∼421.5–440 nm (Figure 2, A-D insets), reflecting direct coordination of the free electron pair of the azole ring nitrogen to the heme iron(43). The maxima and minima of these binding spectral curves were used to plot the binding isotherm and calculate the *K*_d_ values for each ligand (Table 1). The four azole drugs tested demonstrated tight binding affinities for CYP168A1 from high to low: ketoconazole (0.684 ± 0.076 µM), miconazole (0.882 ± 0.182 µM), econazole (2.46 ± 0.42 µM), and clotrimazole (2.99 ± 0.39) (Figure 2, Table 1). Notably, larger azoles exhibited a higher binding affinity (lower *K*_d_) than their corresponding smaller ligand counterparts, with the largest ligand – ketoconazole – having a sub-micromolar *K*_d_ (Table 1). In contrast, binding of various fatty acids to the CYP168A1 ferric heme iron caused a characteristic Type I shift of the Soret band to 380-392 nm, reflecting displacement of the heme distal water at the sixth ligand to the heme iron and indicative of a substrate binding in the active site (Figure 3A-G insets). As done with the Type II azole ligands, the maxima and minima of these binding spectral curves were used to calculate the *K*_d_ values for each fatty acid examined (Figure 3, Table 1). Consistent with our observations of larger azoles binding to the enzyme, higher molecular weight long-chain fatty acids were preferred over their short chain counterparts, with the highest affinity ligands being palmitic (0.207 ± 0.038 μM), stearic (0.327 ± 0.072 μM), and oleic (0.374 ± 0.065 μM) acids (Figure 3, Table 1). Interestingly, while exhibiting a preference for longer chain hydrocarbons, CYP168A1 seemed to bind both monounsaturated and polyunsaturated fatty acids with similar, sub-micromolar affinities (Table 1). The Type I difference spectra and tight (sub-micromolar, in most cases) binding affinities suggested that fatty acid ligands of chain length C10 or greater had the potential to be substrates of CYP168A1. Ligand binding was also assessed for steroids (e.g., cholesterol, etc.) and certain drugs (e.g., raloxifene, ciprofloxacin), but no spectral changes in the Soret bands were observed during the titrations.

**Figure 2.**
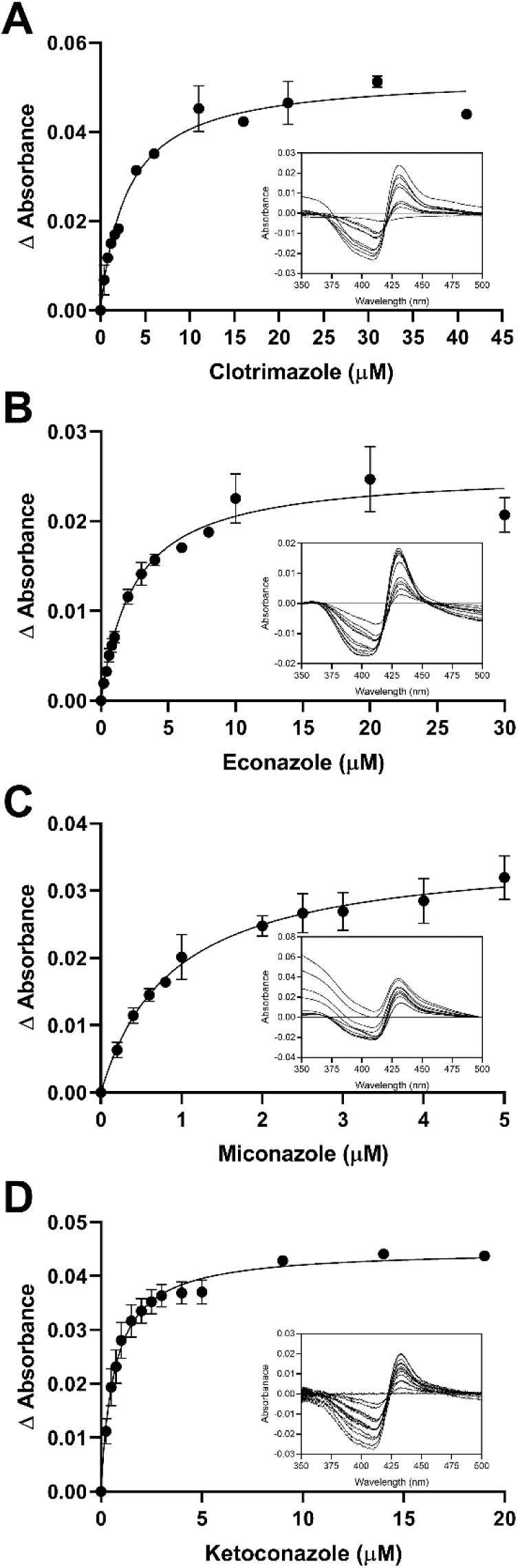
Azole Binding Isotherms with Representative Binding Spectra for CYP168A1. Binding isotherms of clotrimazole (A), econazole (B), miconazole (C) and ketoconazole (D), with insets containing representative binding spectra, were fitted to the one binding site model with R^2^ of 0.937, 0.899, 0.871 and 0.939, respectively.

**Figure 3.**
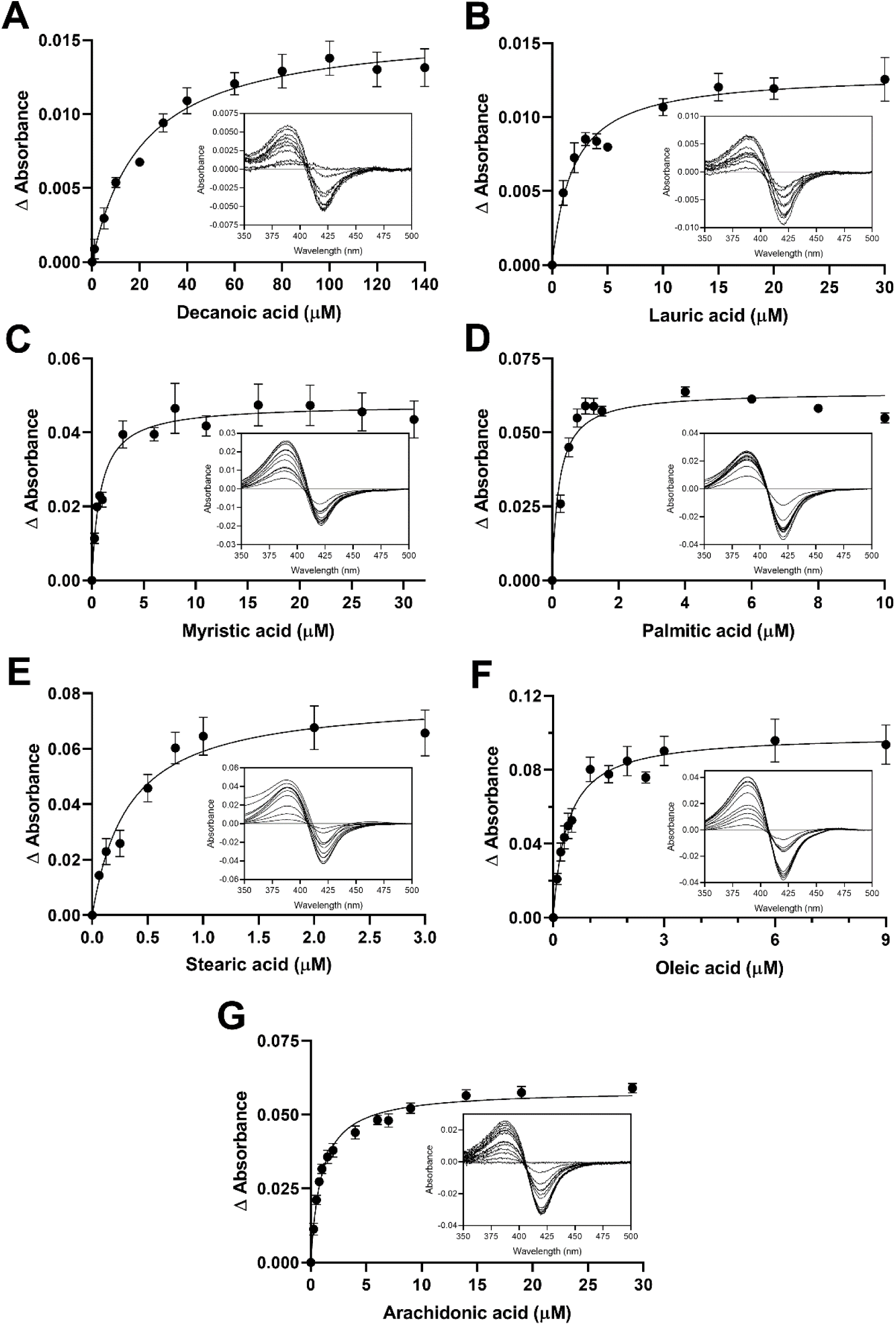
Fatty Acid Binding Isotherms with Representative Binding Spectra for CYP168A1. Binding isotherms of decanoic (A), lauric (B), myristic (C), palmitic (D), stearic (E), oleic (F) and arachidonic (G) acids, with insets containing representative binding spectra, were fitted to the one binding site model with R^2^ of 0.935, 0.906, 0.880, 0.908, 0.897, 0.878 and 0.967, respectively.

**Table 1:** CYP168A1 Binding Constants for Fatty Acids and Azoles.

| Ligands | | $K_d \pm SE$ (μM) | $\Delta$ Absorbance maxima (AU) |
| --- | --- | --- | --- |
| Fatty acids |  |  |  |
| Hexanoic acid | C6:0 | ND | ND |
| Octanoic acid | C8:0 | ND | ND |
| Decanoic acid | C10:0 | $21.0 \pm 3.5$ | 0.0159 |
| Lauric acid | C12:0 | $1.87 \pm 0.33$ | 0.0130 |
| Myristic acid | C14:0 | $0.833 \pm 0.149$ | 0.0476 |
| Palmitic acid | C16:0 | $0.207 \pm 0.038$ | 0.0638 |
| Stearic acid | C18:0 | $0.327 \pm 0.072$ | 0.0784 |
| Oleic acid | C18:1 | $0.374 \pm 0.065$ | 0.0992 |
| Arachidonic acid | C20:4 | $0.960 \pm 0.074$ | 0.0583 |
| Azoles |  |  |  |
| Clotrimazole | | $2.99 \pm 0.39$ | 0.0527 |
| Econazole | | $2.46 \pm 0.42$ | 0.0256 |
| Miconazole | | $0.882 \pm 0.182$ | 0.0359 |
| Ketoconazole | | $0.684 \pm 0.076$ | 0.0449 |

ND: no spectral shift detected; SE: standard error.

### Lauric Acid Metabolism by CYP168A1

The reductase electron transfer partners of CYP168A1 are currently unknown. Therefore, in order to determine if CYP168A1 was able to catalyze the oxidation of the model substrate lauric acid, initial experiments were conducted using either a series of hydroperoxide compounds as oxygen surrogates and electron donors or a spinach-derived redox partner complex consisting of the spinach ferredoxin (Fdx) and ferredoxin reductase (FdR). The spinach Fdx and FdR redox partners were previously shown to function as effective electron transfer surrogates to promote oxidation of substrates by the bacterial CYPs CYP125A1(44) and CYP141(30), structurally related CYP enzymes from *M. tuberculosis*. Gas chromatography-mass spectrometry (GC-MS) experiments confirmed that the major metabolite produced was the 11-hydroxylauric acid (Figure 4). The mass spectrum of the trimethylsilyl-derivatized lauric acid metabolite formed in CYP168A1 incubations with the spinach redox partners (Figure 4B) matched the fragmentation pattern of the derivatized 11-hydroxylauric acid standard (Figure 4A). In addition, the *m/z* fragment of 117, corresponding to the (CH_3_)_3_SiO(CH_2_-CH_3_) ion, is a characteristic fragment of an ω-1 hydroxyl fatty acid derivative(45). Next, liquid chromatography-mass spectrometry (LC-MS) experiments were employed to quantify formation of the 11-hydroxylauric acid metabolite with the hydroperoxide compounds or the spinach redox partners. The 11- and 12- hydroxylauric acid standards were baseline separated (Figure 5). The LC-MS trace for the lauric acid incubation of CYP168A1 with the spinach redox partners and the cofactor nicotinamide adenine dinucleotide phosphate (NADPH) shows the 11-hydroxylauric acid as the major metabolite formed and presence of a minor hydroxyl metabolite matching the retention time of the 12-hydroxylauric acid, although the identity of the minor metabolite could not be confirmed by GC-MS due to its low abundance.

**Figure 4.**
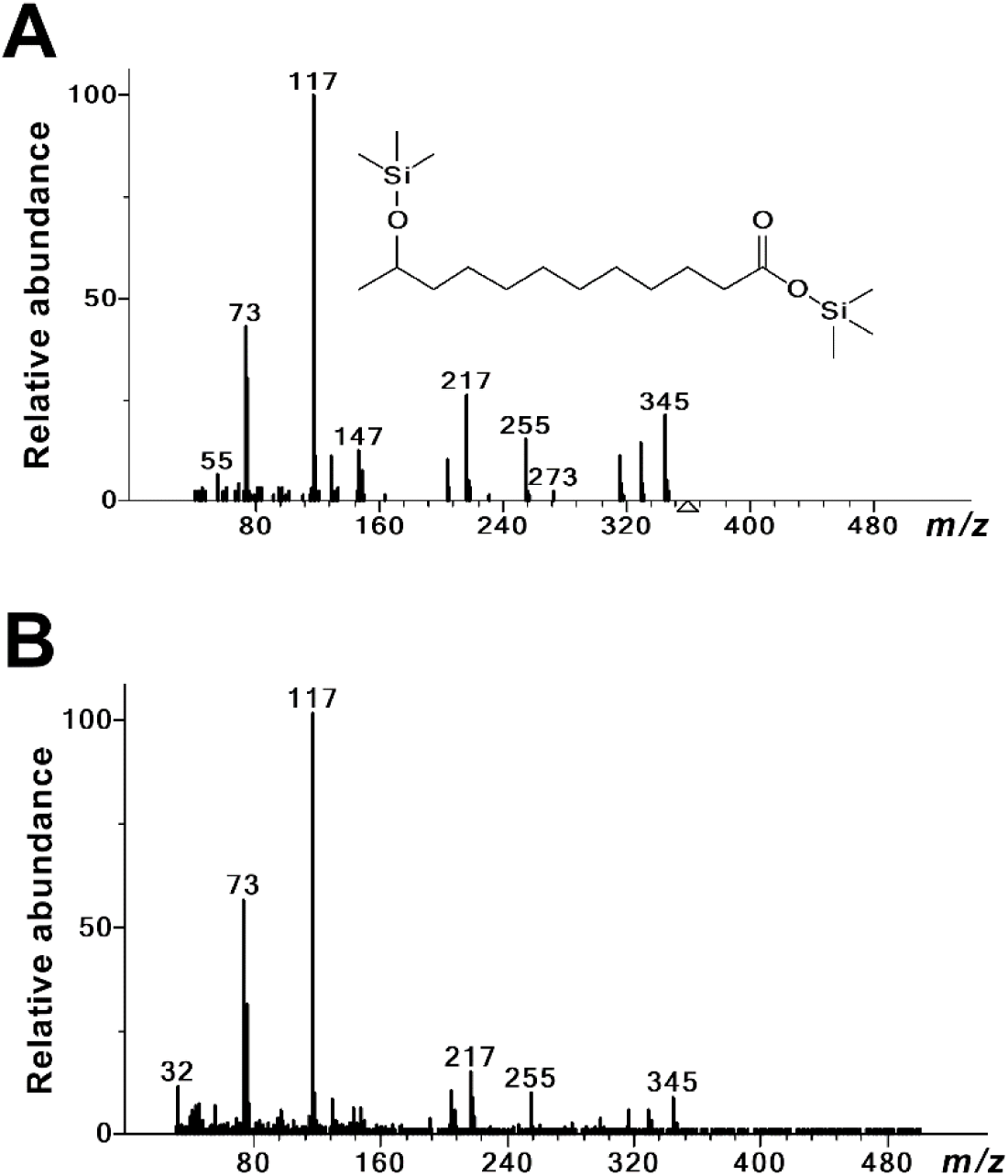
*P. aeruginosa* CYP168A1 Metabolizes Lauric Acid to the 11-Hydroxylauric Acid. Mass spectra of the trimethylsilyl-derivatized 11-hydroxylauric acid standard (A) and CYP168A1 lauric acid metabolite (B).

**Figure 5.**
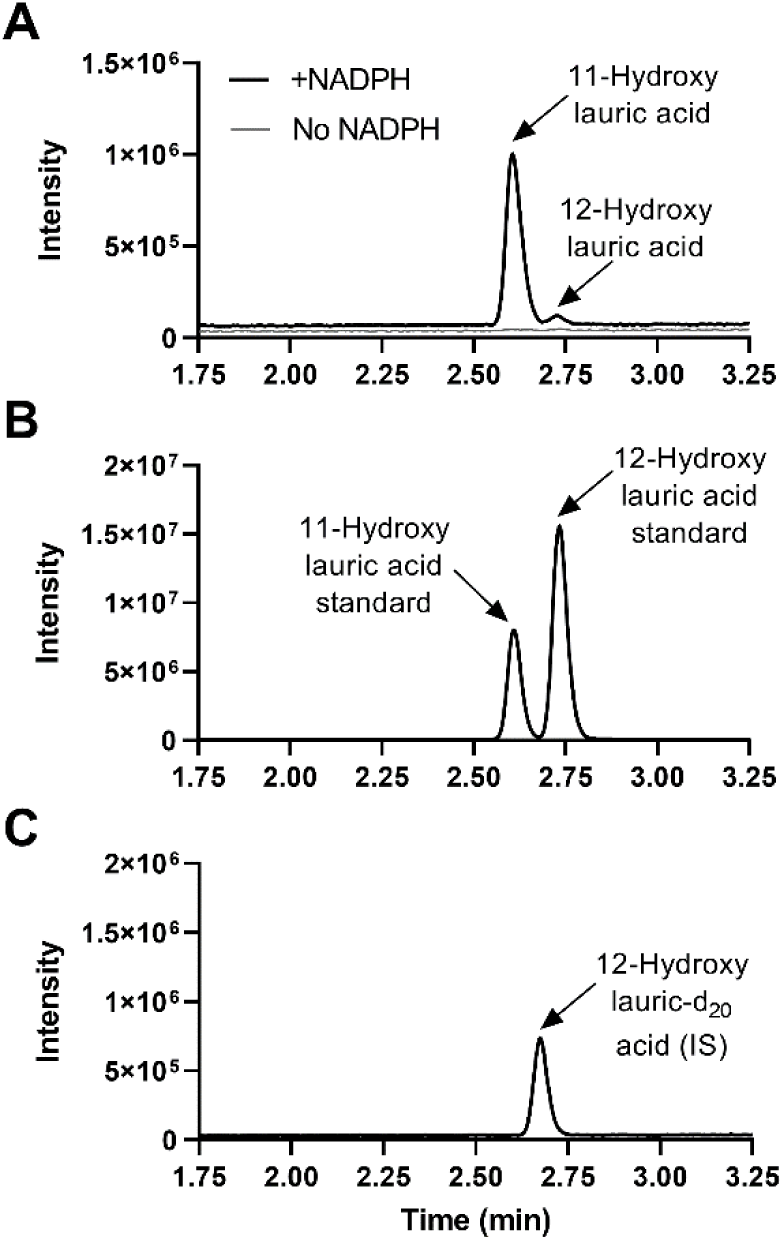
LC-MS Chromatograms of the 11- and 12-Hydroxylauric Acid Metabolites. Representative Multiple Reaction Monitoring (MRM) chromatograms for the hydroxyl lauric acid metabolites formed in incubations of CYP168A1 with lauric acid at 10 µM and in presence or absence of NADPH (A). Representative MRM chromatogram for the 11- and 12-hydroxylauric acid standards at 5 µM (B). Representative MRM chromatogram for the internal standard (IS) 12-hydroxylauric-d_20_ acid (C).

### Hydroperoxide-Driven Catalysis by CYP168A1

A concentration range of the tert-butyl (tBPH) and cumene hydroperoxides (CuOOH) were tested for catalysis of lauric acid hydroxylation by CYP168A1 and formation of the 11-hydroxyl derivative. As can be observed in Figure 6, maximal metabolite formation occurred following incubations of 1 µM CYP168A1 with 10 µM lauric acid for 120 min with the hydroperoxides. The results demonstrate that CYP168A1 was able to hydroxylate lauric acid utilizing the peroxide shunt pathway, bypassing the catalytic cycle necessary for transferring electrons from NAD(P)H through a cytochrome P450 reductase(46). Maximal lauric acid metabolism was achieved when the concentration of each hydroperoxide was 0.25 mM. However, metabolism was uniquely altered for each hydroperoxide in a concentration-dependent manner.

**Figure 6.**
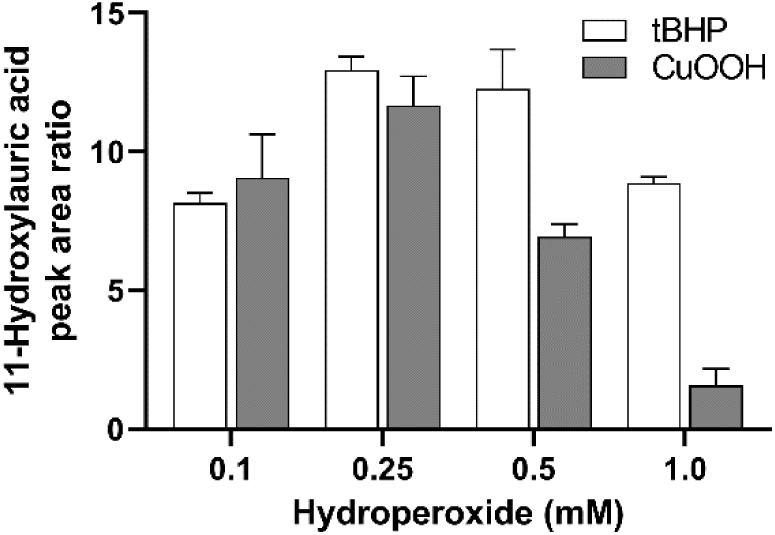
Effect of Hydroperoxides on Lauric Acid Catalysis by CYP168A1. The relative quantification of the 11-hydroxylauric acid metabolite formed by CYP168A1 with the hydroperoxides (tBPH, tert-butyl hydroperoxide and CuOOH, cumene hydroperoxide) was achieved by LC-MS and peak area ratios of the 11-hydroxylauric acid were reported with their respective standard deviations represented graphically as error bars.

### Redox Partner-Driven Catalysis by CYP168A1

In order to investigate ligand metabolism requiring the transfer of electrons along the full P450 catalytic cycle, surrogates for the bacterial reductases were employed. We used the spinach Fdx and FdR as redox partners in the following experiments. The effects of varying Fdx and FdR concentrations were assessed in order to determine the optimal CYP:redox partner ratio (Figure 7). When increasing Fdx concentrations, formation of the 11- hydroxyl metabolite was maximum at the 1/20 CYP:Fdx ratio (Figure 7A). In contrast, when increasing FdR concentrations, formation of the 11-hydroxylauric acid reached plateau at 0.2 U/mL of FdR (Figure 7B). The optimal ratio of CYP:Fdx:FdR was determined to be 1 µM:20 µM:0.2 U/mL for maximal CYP168A1 lauric acid hydroxylation activity. However, under the 1/20 CYP:Fdx ratio conditions and the 0.2 U/mL FdR, metabolism of lauric acid exceeded the 20% substrate consumption leading to non-linear enzyme kinetics. Thus, to allow for further assessment of CYP168A1 lauric acid hydroxylation kinetics under appropriate steady state conditions, we set the final CYP:Fdx:FdR ratio at 1 µM:10 µM:0.05 U/mL.

**Figure 7.**
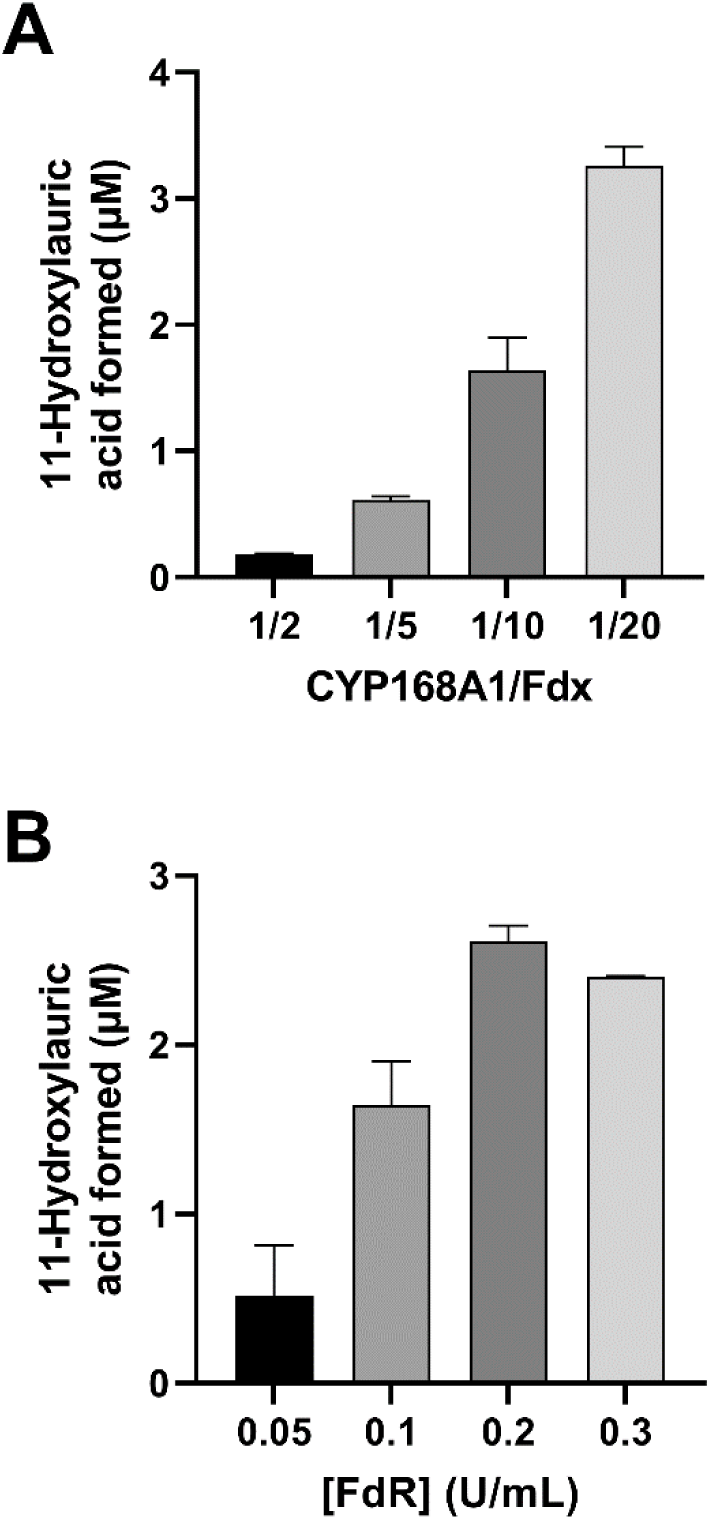
Effect of the Spinach Redox Partners on Lauric Acid Catalysis by CYP168A1. Quantification of the 11-hydroxylauric acid metabolite formed with increasing Fdx concentrations at fixed CYP168A1 (1 µM) and FdR (0.1 U/mL) concentrations (A) and with increasing FdR concentrations at fixed CYP168A1 (1 µM) and Fdx (10 µM) concentrations (B). The bar graphs represent the mean of assays performed in triplicate, with error bars representing standard deviations.

### Kinetics of Lauric Acid ω-1-Hydroxylation by CYP168A1

To further elucidate the kinetic mechanism of lauric acid oxidation by CYP168A1, steady-state kinetic experiments were performed using the optimized reaction conditions described above. Due to the sigmoidal nature of the data (Figure 8), both the Michaelis-Menten and Hill fits were compared using the Akaike Information Criterion (AIC) (47). The difference of the second order Aikake Information Criterion (AICc) is representative of the difference between the simpler model (Michaelis-Menten) minus the alternative model (Hill), which includes more fitting parameters. The difference in AICc for lauric acid ω-1-hydroxylation by CYP168A1 was determined as 16.85 (Table 2). A positive number for the difference of AICc means that the alternative model (with more parameters) has the lower AICc and is preferred. Thus, the Hill model was judged to yield a better fit for the lauric acid kinetic data set with a correct fit probability of 99.98% (Table 2). According to the Hill equation, CYP168A1 has an *S_50_* of 33.7 ± 2.2 µM, a *V_max_* of 0.118 ± 0.004 nmol/min/nmol P450 and an *n* value, a measurement of the cooperativity of substrate binding, of 1.46 (Figure 8, Table 2). An *n* value above 1 is indicative of positive cooperativity in substrate binding(48).

**Figure 8.**
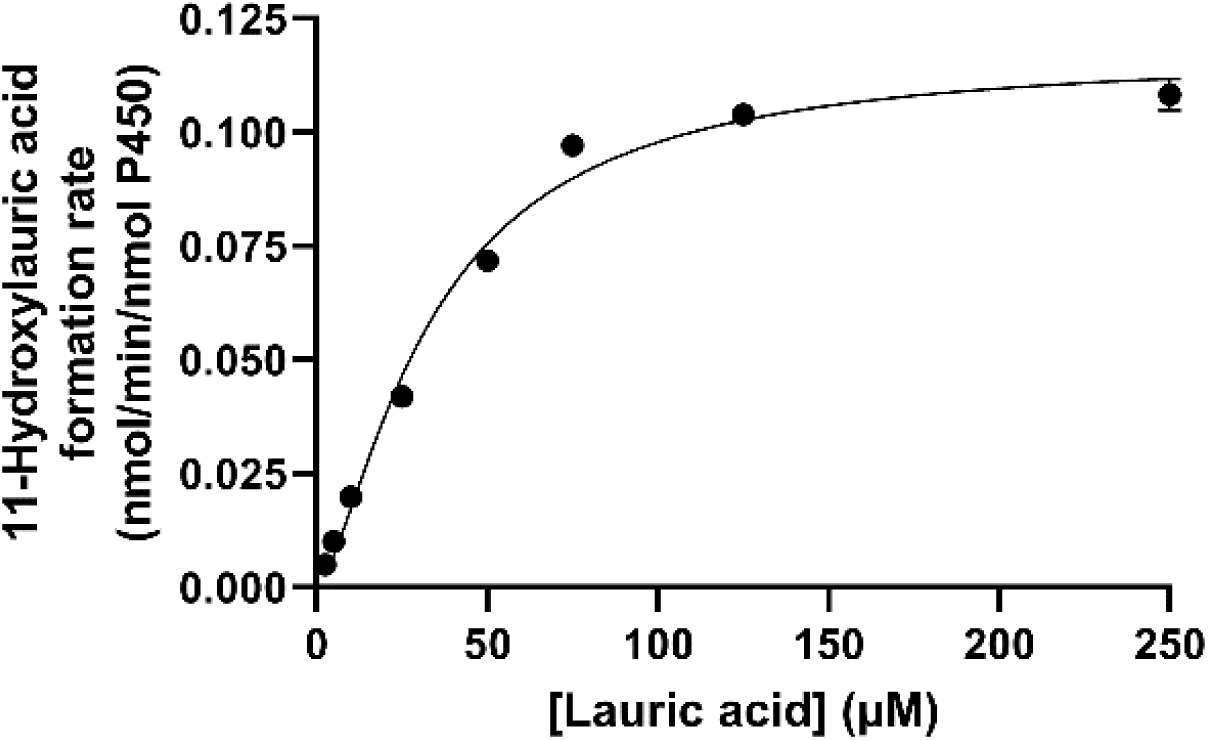
Kinetic of Lauric Acid ω-1-Hydroxylation by CYP168A1. Lauric acid ω-1-hydroxylation by CYP168A1 best fitted the Hill equation. Each data point represents the mean of assays performed in triplicate, with error bars representing the standard deviations (some of the error bars being too small to be observed). The coefficient of determinations, R^2^, for the regression model fit of the 11-hydroxylauric acid kinetic was 0.990.

**Table 2:** Comparison of Michaelis-Menten and Hill Fits for Lauric Acid ω-1-Hydroxylation by CYP168A1.

| Model fit | $K_m/S_{50} \pm \text{SE}$<br>($\mu\text{M}$) | $V_{max} \pm \text{SE}$<br>(nmol/min/nmol P450) | $R^2$ | Correct fit<br>probability | Difference<br>of AICc |
| --- | --- | --- | --- | --- | --- |
| Michaelis-Menten | $46.1 \pm 5.4$ | $0.138 \pm 0.006$ | 0.977 | 0.02% | 16.85 |
| Hill | $33.7 \pm 2.2$<br>($n = 1.46$) | $0.118 \pm 0.004$ | 0.990 | 99.98% | |

$n$ : cooperativity value for the substrate binding to the enzyme, SE: standard error, AICc: second order Aikake Information Criterion.

### Arachidonic Acid Metabolism by CYP168A1

The low *K*_d_ measured for the longer unsaturated fatty acids drove our investigation towards characterizing the potential for arachidonic acid metabolism by CYP168A1. The CYP:redox partner ratio previously optimized for lauric acid kinetic experiments was initially used for linearity assessment of the arachidonic acid metabolism. LC-MS experiments were carried out with arachidonic acid and CYP168A1 where two major hydroxyl metabolites were identified, the 18-hydroxyeicosatetraenoic acid (18-HETE) and the 19-HETE (Figure 9A). Formation of the 20- hydroxyeicosatetraenoic acid (20-HETE) metabolite was also detected, but at too low level to allow quantification (Figure 9A). The MSMS spectra of the 18- (Figure 9C) and 19-HETE (Figure 9B) formed in CYP168A1 arachidonic acid incubations matched the MSMS spectra of the corresponding authentic standards. Quantification of the 18-HETE and 19-HETE formed over time with CYP168A1 (data not shown) highlighted the requirement in optimizing the concentrations of the CYP168A1 enzyme and the spinach redox partners in the arachidonic acid incubations to achieve steady state conditions. Effectively, the CYP:Fdx:FdR ratio had to be adjusted to 0.5 µM:5 µM:0.025 U/mL to achieve linearity in product formation up to 20 min (data not shown). Concurrently to our linearity assessment, we tested the effect of the azole ligand ketoconazole on the 18- and 19-HETE metabolite formation (Figure 10). In presence of 10 µM ketoconazole, CYP168A1 formation of 18- and 19-HETE was inhibited to 67.6% (Figure 10A) and 62.7% (Figure 10B), respectively, compared to the solvent control.

**Figure 9.**
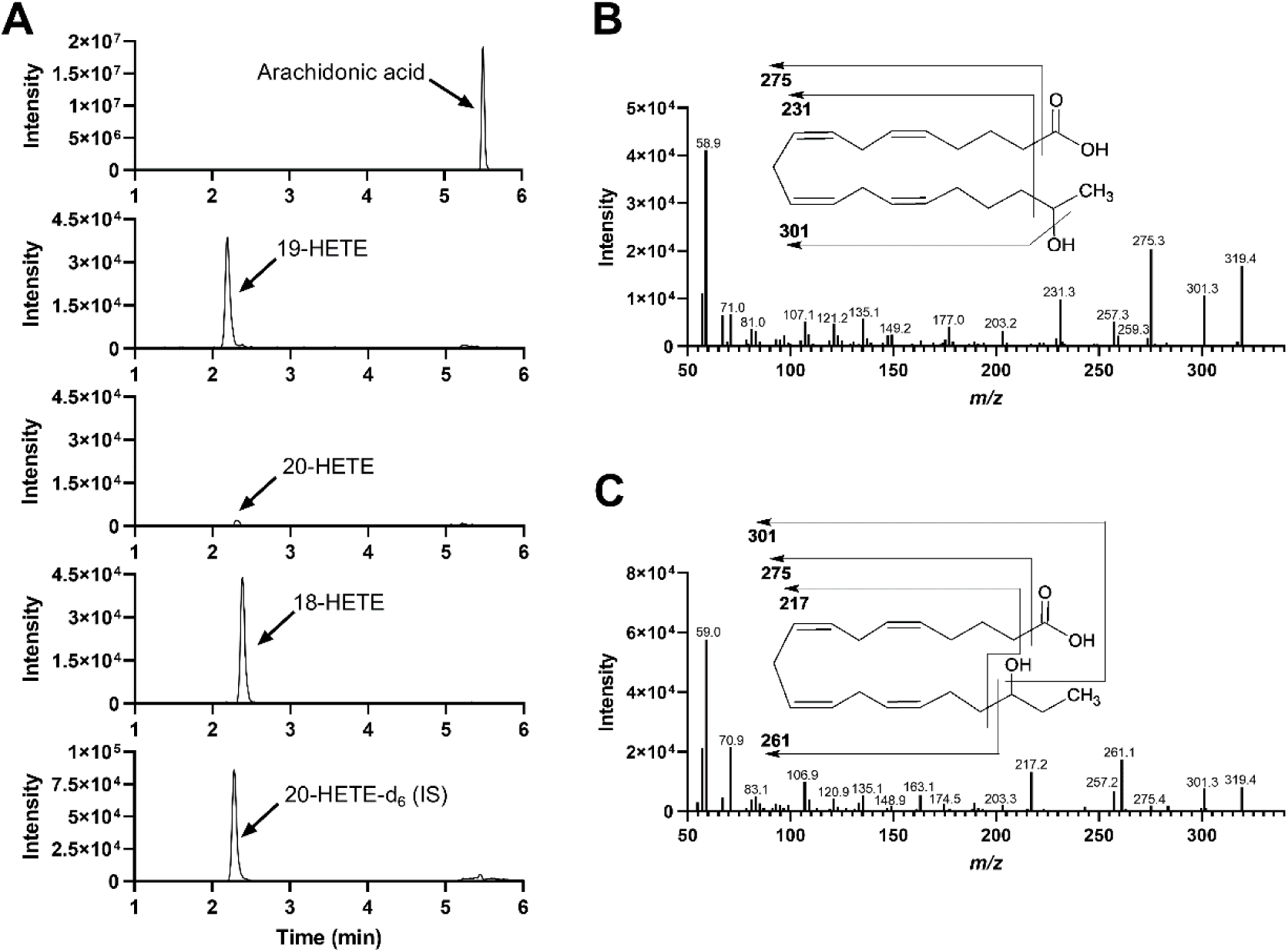
*P. aeruginosa* CYP168A1 Metabolizes Arachidonic Acid to the 18- and 19-HETE Metabolites. Representative MRM chromatograms for the arachidonic acid and its 18-, 19- and 20-HETE metabolites formed in incubations with the recombinant CYP168A1 in presence of the spinach redox partners and the co-factor NADPH, including the MRM chromatogram for the internal standard 20- HETE-d_6_ (A). LC-MS/MS spectra for the 19-HETE (B) and 18-HETE (C) metabolites formed in arachidonic acid incubation with CYP168A1, including structure fragmentation insets.

**Figure 10.**
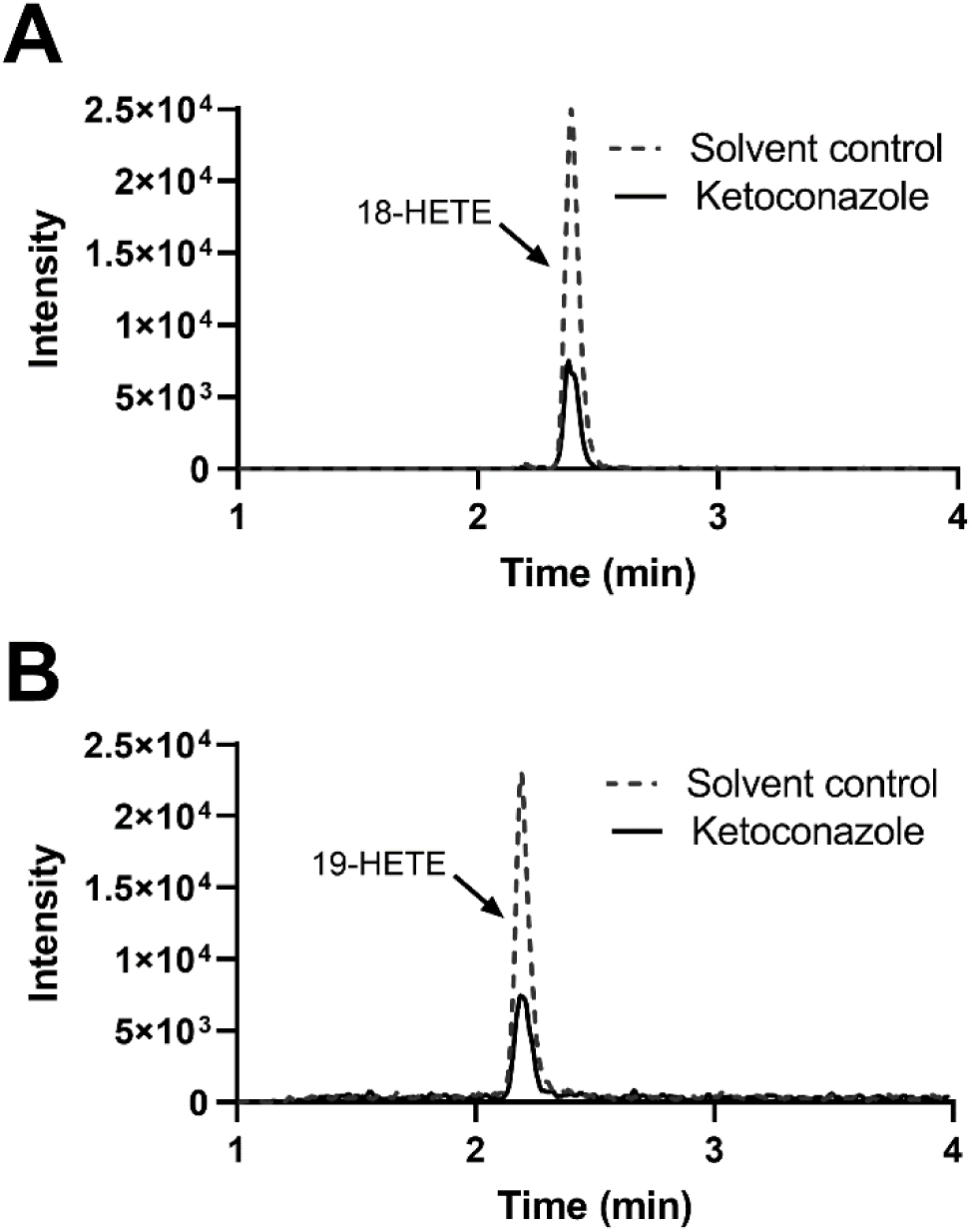
Inhibition by Ketoconazole of CYP168A1 18- and 19-HETE Formation. Representative MRM chromatograms for 18- (A) and 19-HETE (B) formed in incubations of arachidonic acid with CYP168A1 using the spinach redox partners and the co-factor NADPH in absence (solvent control, dotted line) or presence of ketoconazole (10 µM, solid line).

### Kinetics of Arachidonic Acid ω-1- and ω-2-Hydroxylation by CYP168A1

To explore the kinetic mechanisms driving the arachidonic acid hydroxylation by CYP168A1, steady state kinetic experiments were performed using the optimized CYP:redox partner ratio described above for arachidonic acid. Unexpectedly, the 18- and 19-HETE formation rates measured over a range of arachidonic acid concentrations displayed substrate inhibition kinetics (Figure 11). According to the substrate inhibition model fit of the 18- and 19-HETE formation rate data, CYP168A1 exhibits a *K_m_* of 36.3 ± 17.9 µM, *V_max_* of 81.7 ± 32.6 pmol/min/nmol P450, and *K_i_* of 13.4 ± 8.1 µM for 18-HETE formation, and a *K_m_* of 41.1 ± 24.6 µM, *V_max_* of 222 ± 110 pmol/min/nmol P450, and *K_i_* of 15.8 ± 7.9 µM for 19-HETE formation.

**Figure 11.**
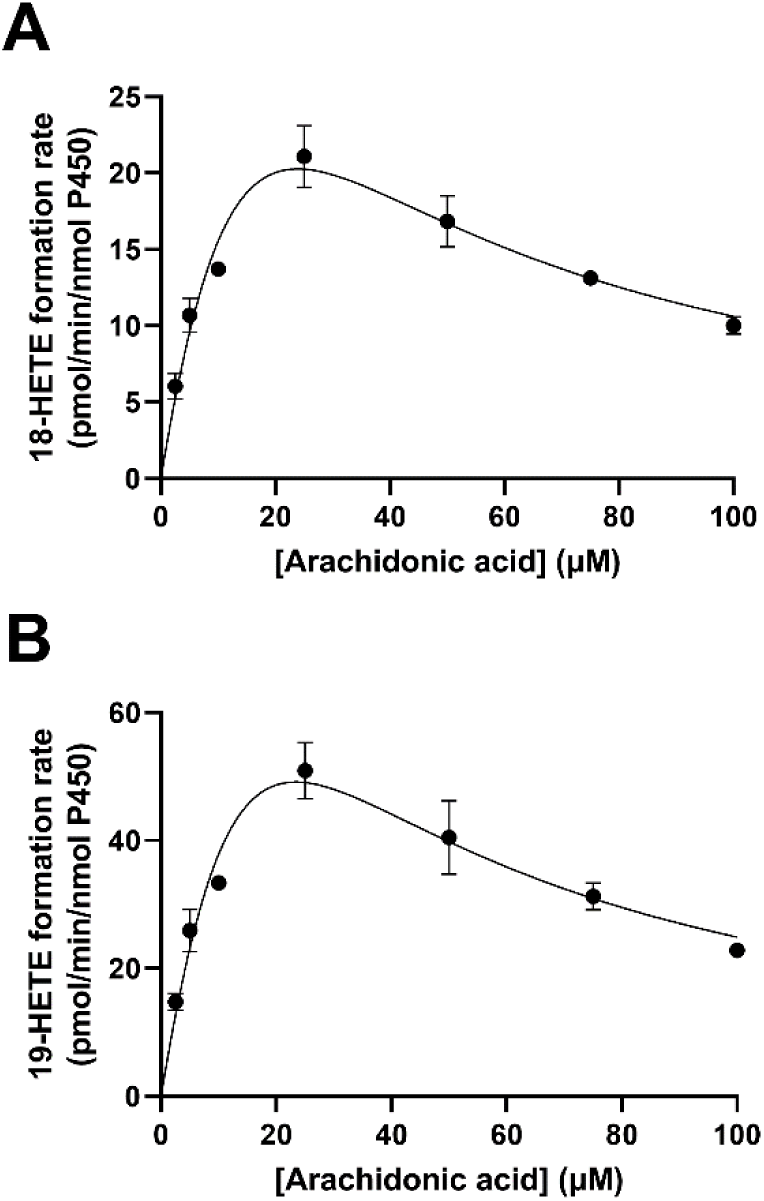
Kinetic of Arachidonic Acid ω-1- and ω-2-Hydroxylation by CYP168A1. Arachidonic acid ω-2- (A) and ω-1-hydroxylation (B) by CYP168A1 fitted the substrate inhibition model. Each data point represents the mean of assays performed in triplicate, with error bars representing standard deviations. The coefficient of determinations, R^2^, for the regression model fit of 18- and 19-HETE kinetics were 0.911 and 0.901, respectively.

### CYP168A1 Homology Model

In an effort to explore the structural basis of fatty acid hydroxylation of CYP168A1, we built a homology model based on CYP VdH, a vitamin D3 (cholecalciferol) hydroxylase from *P. autotrophica*(49) using USCF MODELLER(50) (Figure 12A). The model demonstrated a high-quality alignment with a zDOPE score of -1.08 and a GA341 score of 1. The PyMol castp plugin calculated an active site volume of 668.6 Å^3^ with an area of 569 Å^2^ (Figure 12B). As observed in other CYP enzyme structures, there is the critical active site threonine (T300), which plays a role in oxygen activation, present in the long I-helix over the heme iron. Overall, the structure is relatively compact, reminiscent of other bacterial CYP enzyme structures (23, 51). Interestingly, the structure contains an abundance of aromatic residues, including 20 Phe, 8 Trp, and 8 Tyr (Figure 12C). A number of the Phe residues appear to be participating in pi-pi stacking interactions, which may contribute to the relative stability observed with the protein, as this has also been seen with some thermostable bacterial CYP enzymes(52).

**Figure 12.**
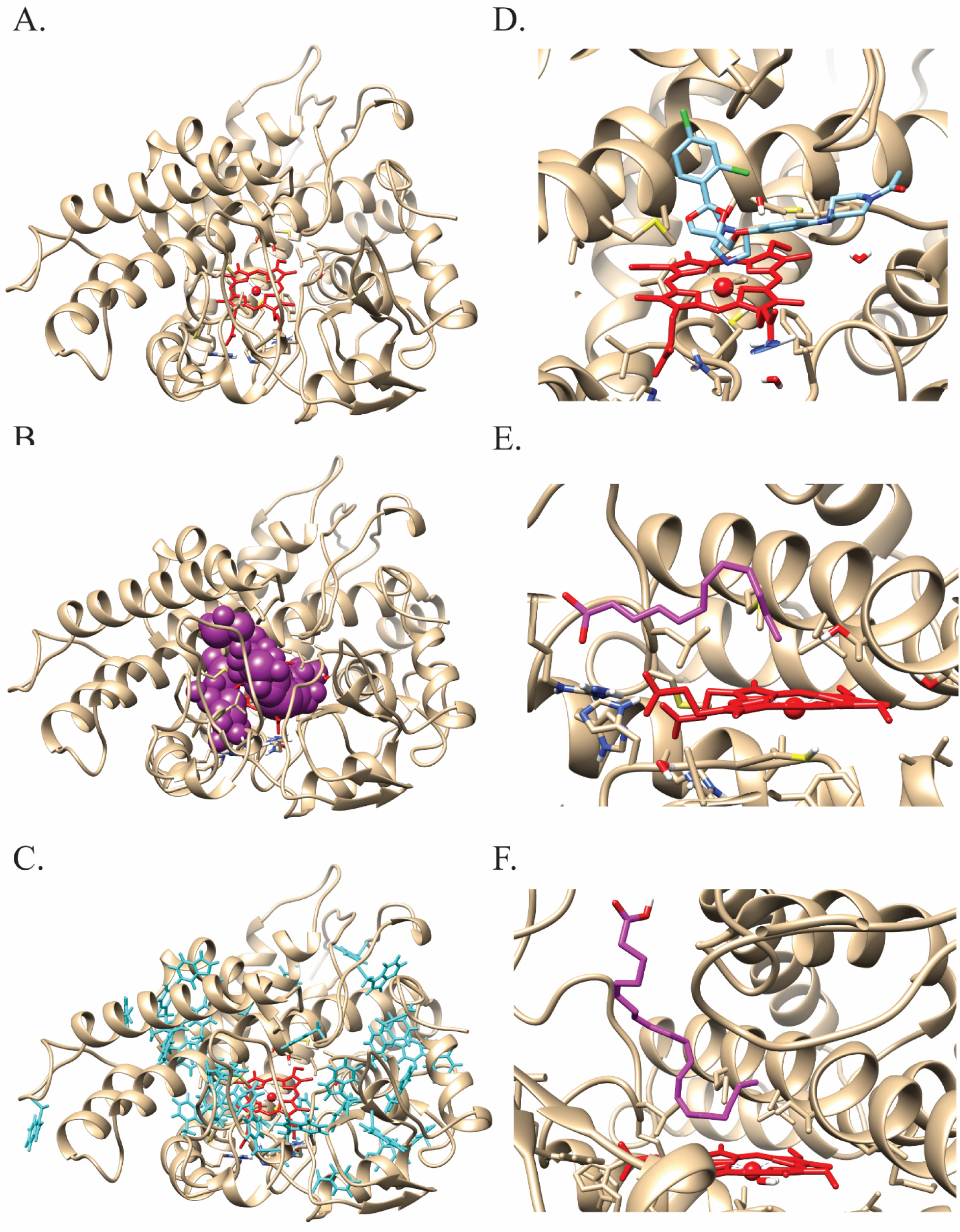
Docking of Ligands to the CYP168A1 Homology Model. (A) Stylized amino acid backbone of the CYP168A1 homology model, with the heme prosthetic group shown in red (in all structures), (B) the P-cast defined active site of the enzyme (represented by magenta balls), (C) aromatic residues of CYP168A1 (shown in cyan), (D) active site cutaway of the docked structure of ketoconazole (cyan) within the CYP168A1 active site, showing the azole nitrogen in close proximity to the heme iron, (E) active site cutaway of docked structure of lauric acid (magenta); for simplicity sake, only the carbon skeleton backbone is shown; note the kinked and extended structure of the substrate, (F) active site cutaway of the docked structure of arachidonic acid (magenta) within the CYP168A1 active site; the conformation of the substrate is similarly kinked and extended as in the case of lauric acid. All figures were generated using UCSF Chimera (https://www.cgl.ucsf.edu/chimera/).

### Docking of Substrates and the Inhibitor Ketoconazole to CYP168A1

To increase our structural understanding of how ligands interact with the enzyme, both substrates (lauric acid and arachidonic acid) and an inhibitor (ketoconazole) were docked to the protein using AutoDock Vina(53). The most thermodynamically stable structure with the docked inhibitor, ketoconazole, with a ΔG of -8.0 kcal/mol, found it present in the active site in a canonical pose with the nitrogen of the azole ring 1.81 Å from the heme iron (Figure 12D). The binding pose is similar to the CYP3A4-ketoconazole structure, with the bulk of the molecule in an extended conformation across the active site, making numerous contacts with hydrophobic residues that line the substrate access channel, including (M292, V343, V445, and V357), which likely help anchor it in a high-affinity inhibitory conformation. While the favored pose for the model substrate lauric acid found it in the active site in close proximity with many of the same hydrophobic residues (including M292, A296, V343, V445, and V357), the carbon backbone of the ligand was kinked at C7, allowing the molecule to bend and place C11 within ∼5 Å of the heme iron (5.02 Å) (Figure 12E). Additional hydrophobic contacts occur with F118, L138, and G346. However, unlike ketoconazole, we also observed contacts with important hydrophilic residues, including the catalytic threonine (T300) and S444, an active site serine residue that has a homology in CYP3A4 and is critical for oxidative activity against a number of substrates, including testosterone(54), carbamazepine(55), and diazepam(56). The most stable binding pose of lauric acid within the CYP168A1 active site returned a binding energy of ΔG of -5.7 kcal/mol. In contrast, the thermodynamically stable binding conformation of arachidonic acid found the molecule in a perpendicular orientation to the heme iron, with the carboxyl moiety extended toward the surface of the enzyme and a kink occurring at C15, positioning C18, C19, and C20 all close to the heme iron (4.719 Å, 5.083, 5.084, respectively) (Figure 12F). The binding energy of the arachidonic acid docked structure was -7.2 kcal/mol, closer to that observed with the inhibitor ketoconazole. All of the same hydrophobic contacts observed with both ketoconazole and lauric acid were also present in the arachidonic acid structure (i.e., F118, L138, V343, G346, M292, V343). However, its larger size permitted additional contacts with residues Q80, L227, A228, M295, and A296. Again, as observed with the lauric acid structure, there was interaction between the substrate and the catalytic threonine (T300).

## DISCUSSION

CYP168A1 is the first cytochrome P450 enzyme from *P. aeruginosa* to be cloned, expressed, and characterized in a heterologous host (42). The genome of *P. aeruginosa* contains four putative CYP enzymes of unknown function (10). CYP168A1 is the largest of these four putative CYP enzymes at 444 amino acids in length, significantly larger than the other three CYPs, particularly given that CYP239A1, at 386 amino acids in length, has been postulated to be a pseudogene (10).

Cytochrome P450 enzymes perform a wide variety of functions in bacteria (51), from secondary metabolite production (20–22) to carbon source metabolism (15–17), to synthesis of lipids critical for cell wall integrity (51, 57). In some cases, these functions are essential to microbial growth (44); therefore, it has been suggested that CYP enzymes may be useful targets for novel antibiotic therapies, particularly for pathogens that are intractable to current treatments (58, 59). *P. aeruginosa* is a gram-negative opportunistic pathogen that is particularly prevalent in the lungs of patients with cystic fibrosis, chronic obstructive pulmonary disease, and nosocomial pneumonia (60–62). In patients with compromised lung and/or immune functions, recurrent infections are common (63, 64). Successive rounds of antibiotic treatment can then lead to resistance, often mediated by the development of biofilms that make it almost impossible to eradicate the organism from the patient’s lung (6, 65). Prior studies have demonstrated that exposure to quinolones and carbapenems, antibiotics commonly used in ICUs, is linked to the development of multidrug-resistant *P. aeruginosa* (66), reducing the therapeutic options available to the clinician and increasing hospital mortality rates. Given this, it is critically important to identify new drug targets to provide effective treatment for these patients.

The UV-visible spectral data obtained from CYP168A1 are consistent with a cytochrome P450 enzyme that is primarily composed of holoprotein, with little P420 species present, indicating that the protein was likely to be active (Figure 1). Initially, in order to determine the variety of ligand chemical space that might be accessible to CYP168A1, we examined several azole drugs, which are known CYP inhibitors (Figure 2) (67). For all the azole drugs tested, a Type II difference spectrum was observed, indicative of inhibitors that bind through coordination of the azole nitrogen free electron pair with the heme iron (68). Interestingly, larger ligands displayed higher affinity with the largest azole tested, ketoconazole, having the highest affinity (0.684 ± 0.076 μM; Table 1). These results are consistent with a rather large and expansive active site, similar to the human CYP3A enzymes (69), that is able to accommodate large, bulky hydrophobic ligands (Figure 12D). Additionally, it points to a strategy for inhibition of the enzyme through a specific azole functionality.

The most closely related CYP enzymes to CYP168A1 that have been thoroughly characterized are from the pathogen *M. tuberculosis* (23,27,67,70). Previous studies have demonstrated that, in the case of this obligate pathogen, CYP121A1 (71, 72), CYP125A1 (44, 73), and CYP142A1(74) are essential for bacterial survival, making them attractive drug targets (58, 75). CYP125A1 and CYP142A1 from *M. tuberculosis* have been identified as cholesterol hydroxylases, suggesting that at least some CYP enzymes from pathogens have evolved functions as steroid or fatty acid hydroxylases, which are common functions for a number of microbial CYP enzymes (76, 77). This knowledge underlined our strategy for identifying possible substrates of CYP168A1 in order to determine its function.

Initially, we examined saturated fatty acids with chain lengths of 6 to 18 carbons (Figure 3; Table 1). As can be observed from the results presented in Figure 3, all fatty acid ligands elicited a Type I difference spectrum indicative of substrate binding, except for those of length C-8 or shorter, where no spectral perturbations were visible. Similar to the trend observed with the azole compounds, CYP168A1 showed a clear preference for longer chain fatty acids, as determined by their *K*_d_, with an optimal length of approximately 16 carbons (Table 1). This suggests a large, or at least deep, hydrophobic active site that can accommodate endogenous long-chain fatty acid substrates that may be present in the host environment (8). Indeed, CYP168A1’s active site volume is quite comparable to the active site volume of CYP3A4 (520 Å^3^) (78) and CYPBM3 (400 Å^3^) (79) (Figure 12B).

Furthermore, CYP168A1 tightly bound both the monounsaturated fatty acid oleic acid (0.374 ± 0.065 μM) and the polyunsaturated fatty acid, arachidonic acid (0.960 ± 0.074 μM), suggesting that both saturated and unsaturated fatty acids could serve as substrates for CYP168A1. Interestingly, some common drugs (e.g., ciprofloxacin and raloxifene) did not exert any changes in the heme Soret spectrum, nor did the steroid cholesterol. This implies that CYP168A1 may have a narrower substrate specificity, limited to fatty acids or similar structurally related molecules.

To determine if fatty acids could indeed be oxidized by CYP168A1, we initially examined catalysis of the model fatty acid substrate, lauric acid, using either the oxygen surrogates of tert-butyl and cumene hydroperoxides or the spinach redox partners. While CYP enzymes are readily identifiable in various bacterial genomes through their unique sequences, such as the FxxGxxxCxG heme motif (10,80,81), it remains a challenge to identify their active redox partners, a ferredoxin and ferredoxin reductase for a typical Type I system (68). Hence, spinach ferredoxin (Fdx) and ferredoxin reductase (FdR) have been employed as an electron delivery system in order to catalyze substrate oxidation for various bacterial CYP enzymes (30, 44). Our initial GC-MS (Figure 4) and LC-MS (Figure 5) experiments with the spinach redox partners confirmed the 11-hydroxylauric acid as the primary lauric acid metabolite formed, validating the identity of CYP168A1 as a sub-terminal fatty acid hydroxylase. In addition, according to our LC-MS analysis, a small proportion (<5%) of lauric acid was metabolized to the ω-hydroxyl derivative (Figure 5A). It is interesting to note that our docking studies determined that lauric acid was present in the active site in a somewhat unusual sterically strained conformation (Figure 12D), which may help explain why CYP168A1 primarily performs the ω-1 oxidation over the ω. After confirming lauric acid metabolite identity, we further examined the tert-butyl (tBHP) and cumene hydroperoxide (CuOOH) concentration-dependency to produce the 11-hydroxylauric acid. Both peroxide compounds equivalently stimulated catalysis of lauric acid with the optimal concentration for each peroxide used being 0.25 mM (Figure 6), a value similar to what has been determined for other CYP enzymes exploiting the peroxide shunt pathway (46). We then sought to identify the optimal ratio of the spinach Fdx and FdR redox partners based on the hydroxylation of lauric acid to 11-hydroxylauric acid (Figure 7). While the effect of Fdx concentration on the rate of 11-hydroxylauric acid was not saturable in our system (Figure 7A), the concentration of FdR achieved maximal formation of metabolite at a concentration of 0.2 U/mL, indicating that FdR was the rate limiting reactant (Figure 7B). After adjusting the ratio of redox partners to CYP enzyme to maintain steady state conditions, we conducted a complete kinetic characterization of the lauric acid substrate oxidation by CYP168A1, including determining both the *K*_m_ and *V*_max_ values for hydroxylation to the 11-hydroxylauric acid product (Figure 8). Somewhat surprisingly, the data best fit to the Hill equation, with a *n* value of 1.46, *S_50_* of 33.7 ± 2.2 µM, and a *V_max_* of 0.118 ± 0.004 nmol/min/nmol P450, suggesting a degree of cooperativity in the oxidation of lauric acid. It is not unusual for CYP enzymes to exhibit cooperativity in substrate oxidation, as this has been observed with a number of bacterial (13,82,83) and mammalian (84–86) enzymes. However, this may be the first report of cooperative oxidation of a substrate by a CYP enzyme from a known human pathogen. While the oxidation of lauric acid defines CYP168A1 as a fatty acid hydroxylase, lauric acid is primarily a model substrate with no specific biological activity. In contrast, arachidonic acid, a tight binding ligand of CYP168A1 (*K*_d_=0.960 μM), is a well-known lipid second messenger involved in the response of cell signaling enzymes and, importantly for *P. aeruginosa* infection, a key regulator of inflammation (87). CYP168A1 metabolized arachidonic acid to predominantly the ω-1 19-HETE and the ω-2 18-HETE products (Figure 9A), as confirmed by their MSMS spectra (Figure 9B,C) matching the authentic HETE standards. It was also able to form the 20-HETE metabolite to a lesser extent (Figure 9A). In order to determine the potential for azole compounds to inhibit this reaction, we conducted the incubation in the presence of 10 µM ketoconazole, which significantly reduced the formation of both metabolites (Figure 10), indicating that ketoconazole is an effective *in vitro* inhibitor of the CYP168A1 mediated oxidation of arachidonic acid. A full kinetic analysis of arachidonic acid metabolism retrieved a *K*_m_ of 36.3 ± 17.9 µM and a *V_max_* of 81.7 ± 32.6 pmol/min/nmol P450 for formation of the 18-HETE metabolite and a *K*_m_ of 41.1 ± 24.6 µM and *V_max_* of 222 ± 110 pmol/min/nmol for formation of the 19-HETE (Figure 11). The production of both 19-HETE and 18-HETE were subject to substrate inhibition, with negligible levels of metabolite being formed at substrate concentrations over 200 μM, and a *K_i_* of 15.8 ± 7.9 µM and 13.4 ± 8.1 µM for 18- and 19-HETE, respectively. Thus, in comparison to the model substrate, lauric acid, which exhibited cooperativity in substrate oxidation, arachidonic acid demonstrates substrate inhibition. In regards to their respective binding conformations in the CYP168A1 active site, arachidonic acid extends from near the heme iron to the roof of the active site cavity, significantly further than the more compact lauric acid (Figure 12E,F). It is likely that the larger size of arachidonic acid in comparison to lauric acid (C20 vs. C12) necessitates it being extended out of the active site, as it is less energetically favorable for it to adopt a more compact conformation within the active site cavity itself. The extended conformation may allow multiple ligands to bind at high concentrations of substrate, thereby giving rise to the observed substrate inhibition.

Recently there has been renewed focus on the physiological relevance of substrate inhibition in enzymatic systems (88, 89). It’s been estimated that up to 20% of all enzyme systems may be subject to substrate inhibition (89). Far from being simply a kinetic artifact, substrate inhibition can have important physiological consequences, including but not limited to: controlling product formation, reducing buildup of toxic metabolic intermediates, or rapid termination of the signal transduction cascade (89). In terms of CYP enzymes, substrate inhibition is a well-known phenomenon that occurs often with the mammalian drug metabolizing CYPs (90). While the biological significance of the substrate inhibition observed with arachidonic acid remains unclear at this juncture, it is interesting to note that it was only detected with arachidonic acid, an inflammatory mediator occurring in the natural environment of the lung, and not with the model substrate lauric acid.

Curiously, arachidonic acid is overproduced in CF patients, including those infected with *P. aeruginosa*, due to a metabolic defect (36, 91). Therefore, this is a common fatty acid that the organism is likely to come into contact with in the lung environment of the CF patient. Indeed, a number of studies have demonstrated the importance of arachidonic acid to the growth and/or virulence of *P. aeruginosa*. Rao et al. established that addition of arachidonic acid and its oxidation products to *P. aeruginosa* cells expressing the metabolic regulator *rahU* led to increased biofilm formation (92). Additional work has shown that an increase in exposure of *P. aeruginosa* to arachidonic acid caused an 8-fold increase in the minimum inhibitory concentration (MIC) for the antibiotic polymyxin B (93), demonstrating a link between arachidonic acid and *P. aeruginosa* antibiotic resistance. Furthermore, this same study revealed that *P. aeruginosa* will incorporate arachidonic acid into its cellular membrane when exposed to it *in vitro* (93), even though arachidonic acid is not synthesized natively by *P. aeruginosa*. In *in vivo* experiments using a rat model of *P. aeruginosa* pneumonia, it was observed that several oxidative metabolites of arachidonic acid, including 20-HETE, contributed to pulmonary vascular hypo reactivity (94). Finally, Auvin and colleagues showed that dietary supplementation of arachidonic acid in a mouse model of *P. aeruginosa* infection led to an increased mortality rate (95). The results from all of these studies point to an important interaction between *P. aeruginosa* and arachidonic acid and its metabolites.

Despite this, little is known of arachidonic acid metabolic pathways in *P. aeruginosa*. To date, only a single enzyme from *P. aeruginosa*, a secreted lipoxygenase known as LoxA, has been identified that is capable of metabolizing arachidonic acid, in this case to 15-HETE (38). Intriguingly, an early study examining metabolism of arachidonic acid derived from human blood polymorphonuclear leukocytes by *P. aeruginosa* produced two oxidative metabolites that were never completely characterized, but whose production was inhibited by carbon monoxide and ketoconazole (37), hallmarks of CYP metabolism and consistent with the results obtained in our study.

While it is well known that CYP metabolites of arachidonic acid, including the EETs and HETEs, are not just metabolic oxidation by-products but are also important regulators of both physiological and pathophysiological processes (34,96,97), their production has never previously been linked to a microbial pathogen CYP enzyme. Due to the hydrophobic nature of the arachidonic acid metabolites, they often tend to accumulate with tissue lipids (98), but upon stimulation with hormones the metabolites are released to act either through paracrine or autocrine pathways. *In vitro* experiments have demonstrated that multiple mammalian CYP enzymes, including CYP2E1, CYP2J9, and CYP2U1, can metabolize arachidonic acid to both the 19- and 20-HETE oxidation products (99–101). In mammals, 19-HETE acts as a potent vasodilator, renal Na^+^-K^+^ ATPase activator, and platelet aggregation inhibitor (102–104). Whereas 20-HETE, the minor CYP168A1 metabolic product, is known to be important in the regulation of vascular tone and blood flow, as well as playing a role in inflammation by stimulating the production of various proinflammatory mediators, including: PGE2, cytokine tissue necrosis factor alpha (TNFα), and the chemokines IL-8, IL-12, IL-14 (105). The physiological role of 18-HETE is less well defined, but it may also be important in the regulation of blood flow and blood vessel contractility (106).

*P. aeruginosa* is an opportunistic pathogen and can exist as an innocuous soil microorganism in the natural environment (107). Indeed, recent work examining the potential of *P. aeruginosa* as a bioremediation vector has demonstrated that its complement of CYP enzymes are capable of oxidizing medium to short chain alkanes (32). It is possible that the *P. aeruginosa* CYPs may have originally evolved to utilize carbon sources readily available in the local environment, such as lauric acid.

Only more recently, evolutionary speaking, has *P. aeruginosa* evolved the ability to adapt to the mammalian lung (108). As a consequence, its complement of enzymes may now be adjusting to new roles. In general terms, the adaption of *P. aeruginosa* CYP168A1 to metabolize arachidonic acid to 18-, 19-, and nominally 20-HETE may reflect a burgeoning ability of the pathogen to modulate the immune response of the host organism in order to make the lung environment more palatable for colonization. A hallmark of pathogen success as a parasite is the ability for it be able to “communicate” with the host organism through the production of proteins and small molecule metabolites that have the ability to modulate the immune response and/or improve the characteristics of the host environment in order to permit pathogen growth and replication (109, 110). *P. aeruginosa* may accomplish this through its metabolism of arachidonic acid to metabolites, such as 18-, 19,- and 20-HETE, that provide important physiological functions. While the exact role of the production of these metabolites by *P. aeruginosa* remains a mystery, it is an area of active investigation in our laboratory, and it may point toward a critical role for *P. aeruginosa* CYP168A1 in the maintenance of infection.

In summary, we have characterized the first CYP enzyme from *P. aeruginosa* as a fatty acid hydroxylase capable of metabolizing arachidonic acid to 18-, 19-, and nominally 20-HETE, all important physiological mediators. Further investigation of the role that this enzyme plays in the pathogen life cycle will likely reveal new insights on its ability to grow and replicate in the mammalian lung and also new potential drug targets.

## EXPERIMENTAL PROCEDURES

### Materials

Lauric, decanoic and stearic acids, and 5-aminolevulinic acid hydrochloride were purchased from Acros Organics (Fair Lawn, NJ). The 11-hydroxylauric and 12-hydroxylauric-d_20_ acid standards were purchased from Santa Cruz Biotechnology. The 12-hydroxylauric, palmitic and arachidonic (oil) acids, clotrimazole, imidazole, tert-butyl (tBPH) and cumene hydroperoxides (CuOOH) were obtained from Sigma-Aldrich (St. Louis, MO). Myristic acid and econazole were purchased at VWR International (Radnor, PA) and ketoconazole was from Toronto Research Chemicals (Toronto, ON, Canada). Oleic acid and miconazole were obtained from Thermo Fisher Scientific (Waltham, MA). Ampicillin, arachidonic acid sodium salt, 18-HETE, 19(S)-HETE, 20-HETE and 20-HETE-d_6_ were all purchased from Cayman Chemical (Ann Arbor, MI). Isopropyl-β-D-1-thiogalactopyranoside (IPTG), phenylmethanesulfonylfluoride (PMSF), glucose-6-phosphate and β-nicotinamide adenine dinucleotide phosphate (NADP^+^) were obtained from Alfa Aesar (Haverhill, MA). Glucose-6-phosphate dehydrogenase, the spinach Fdx and FdR were purchased from Sigma-Aldrich (St. Louis, MO). All other chemicals and solvents were obtained from standard suppliers and were of reagent or analytical grade.

### Construction of CYP168A1 Expression Vector and Expression of the Recombinant CYP168A1 Protein

The National Center for Biotechnology Information (NCBI) amino acid sequence NP_251165 is a 444 amino acid sequence classified as a putative cytochrome P450 from *P. aeruginosa* PAO1 (111) and designated according to the P450 nomenclature as CYP168A1 (10). The amino acid sequence was initially reverse translated to DNA using the Sequence Manipulation Suite website (112). CYP168A1 cDNA sequence was then codon optimized for expression in *E. coli* using the GenScript GenSmart Codon Optimization Tool (https://www.genscript.com/gensmart-free-gene-codon-optimization.html). Finally, the codon optimized DNA sequence was engineered with four histidine residues at the 3’-end of the sequence prior to the stop codon and inserted into a pUC57 vector using *Nde*I and *Hind*III engineered restriction site sequences at the 5’ and 3’ ends of the CYP168A1 DNA coding sequence, respectively. Following transformation of *E. coli*-DH5α cells (Invitrogen, Carlsbad, CA), pUC57-CYP168A1 plasmids were isolated using the Qiagen Miniprep Kit (Qiagen, Hilden Germany). The CYP168A1 optimized cDNA insert was removed from the plasmid using *Nde*I and *Hind*III restriction enzymes and isolated using agarose gel electrophoresis and the Qiagen Gel Extraction Kit (Qiagen) for ligation into a similarly digested and isolated pCWOri^+^ CYP expression vector (113). This plasmid, designated as pCWOri-CYP168A1, was then used to transform *E. coli*-DH5α cells in preparation for expression of the CYP168A1 protein.

CYP168A1 was expressed under the control of the *tac* promoter of the pCWOri^+^ plasmid using *E. coli*-DH5α cells in Terrific Broth (TB) medium. Briefly, 10 mL of an overnight pCWori-CYP168A1 *E. coli*-DH5α starter culture consisting of Luria-Bertani medium supplemented with 200 µg/mL ampicillin was used to inoculate each liter of TB (also containing 200 µg/mL ampicillin). The bacterial culture was incubated at 37°C under agitation (250 rpm) until the optical density reached an absorbance at 600 nm of 0.5 to 0.8. Then IPTG and 5-aminolevulinic acid were added at 0.5 and 0.25 mM, respectively. The expression culture was allowed to grow for another 24 h at 25°C and 180 rpm agitation. The bacterial cells were then pelleted by centrifugation at 3,400 × *g* and 4°C for 40 min and the cell pellets were stored at -80°C until purification of the expressed CYP168A1 enzyme.

### Purification of the (His)-Tagged CYP168A1

Expressed CYP168A1 was purified using fast protein liquid chromatography with a HisTrap-HP affinity column (GE Healthcare, Chicago, IL). The bacterial cell pellets were thawed on ice and resuspended in buffer A consisting of 50 mM Tris-HCl pH 7.5, 50 mM NaCl, 0.1 mM ethylenediaminetetraacetic acid (EDTA), 20 mM imidazole, and 1 mM PMSF. About 4 mL of buffer A was used for resuspension of a gram of bacterial cell pellet. After addition of lysozyme (0.3 mg/mL) and DNase (700 U), the bacterial cell suspension was stirred on ice for 30 min. Cells were then lysed on ice using a Branson sonicator set at 50% power and three 4-min bursts with 2 min resting time between each burst. Following cell lysis, whole cells and cell debris were separated by ultracentrifugation at 100,000 × *g* and 4°C for 60 min.

The ultracentrifugation supernatant containing the recombinant CYP168A1 protein was loaded onto a HisTrap-HP column (5 mL) previously equilibrated with buffer A. The column was subsequently washed with 5 column volumes of buffer A. CYP168A1 was then eluted with a gradient of imidazole using the elution buffer B (50 mM Tris-HCl, 0.1 mM EDTA, 0.2 M imidazole). Red-colored fractions were analyzed for purity by SDS-PAGE gel electrophoresis and fractions containing the bulk of the recombinant CYP168A1 protein were pooled and then dialyzed at 4°C in 50 mM Tris-HCl pH 7.5, 0.1 mM EDTA, and 0.1 mM DTT. The protein concentration of the purified CYP168A1 was determined by bicinchoninic acid (BCA) assay (Pierce, Thermo Fisher Scientific, Waltham, MA) and the final concentration of the ferrous-CO protein was determined using UV-visible spectroscopy, with an extinction coefficient of є = 91 mM^-1^.cm^-1^ at the wavelength of 450 nm (114). UV-visible spectroscopy was further used to characterize the spectral absorption pattern of the absolute and reduced CYP168A1 protein.

### Ligand K_d_ Determination by Optical Difference Spectroscopy

To determine ligand selectivity of CYP168A1, UV-visible difference spectra were acquired on a Varian Cary 50 Bio UV-visible scanning spectrophotometer (Agilent, Santa Clara, CA) for various ligands, including antifungal azole compounds and fatty acids. Both sample and reference chambers contained 1 mL of 1 µM CYP168A1 in 100 mM potassium phosphate, pH 7.4. Prior to initiating the titration, a baseline was recorded between 350 and 500 nm. Aliquots of ligand stock solutions prepared by serial dilution in dimethyl sulfoxide (DMSO) were added to the sample cuvette, whereas equal volume of vehicle solvent was added to the reference cuvette to determine the difference spectrum at varying concentrations. The absolute changes in absorbance deriving from a minimum of triplicate titrations were plotted as a function of ligand concentration and fitted to the one binding site model using the GraphPad Prism software (version 9.0.0, GraphPad software, La Jolla, CA).

### Recombinant CYP168A1 Lauric Acid in Vitro Metabolic Assays

Due to its high affinity for CYP168A1 and known activity as a substrate for other microbial CYP enzymes (52, 77), lauric acid was used as a model substrate for metabolism studies. For the hydroperoxide-driven catalysis, solutions of tBPH and CuOOH were freshly prepared in DMSO and incubated at different concentrations up to 120 min with 1 µM CYP168A1 and 10 µM lauric acid in 100 mM potassium phosphate, pH 7.4 (46). Reactions were stopped by the addition of an equal volume of ice-cold methanol containing 60 ng/mL 12-hydroxylauric-d_20_ acid as internal standard. Samples were centrifuged at 2,500 × *g* and 4°C for 20 min for protein precipitation. Supernatants were transferred to high performance liquid chromatography (HPLC) vials, and aliquots of 5 µL were analyzed by LC-MS. For the redox partner-driven catalysis, different concentrations of spinach Fdx and FdR were assessed to obtain the optimal CYP/redox partner ratio. Initial linearity experiments were done establishing linearity up to 30 min. The incubations with the spinach redox partners (200 µL) were carried out in 100 mM potassium phosphate, pH 7.4 and 3 mM MgCl_2_ with 1 µM CYP168A1 and 10 µM lauric acid. Concentrations of spinach Fdx and FdR were between 2 and 20 µM and 0.05 to 0.3 U/mL respectively. After an equilibration at 37°C for 3 min, the reactions, prepared in triplicate, were initiated by the addition of a NADPH-regenerating system mix consisting of NADP^+^ (1 mM), D-glucose-6-phosphate (10 mM) and glucose-6-phosphate dehydrogenase (2 IU/mL). The reactions were incubated for 30 min at 37°C under agitation and were stopped by the addition of ice-cold methanol (200 µL) containing 60 ng/mL 12-hydroxylauric-d_20_ acid internal standard. Incubations without the NADPH-regenerating system mix served as negative controls. Precipitated proteins were collected by centrifugation of the stopped reaction samples for 20 min at 2,500 × *g* and 4 °C. Supernatants were transferred to HPLC vials, and aliquots of 5 µL were analyzed by LC-MS. The 11-hydroxylauric acid metabolite was quantified based on a calibration curve ranging from 0.1 µM to 10 µM.

For the kinetic reactions, similar assay conditions were used with concentrations of lauric acid ranging from 2.5 to 250 µM. To ensure steady state kinetic conditions and less than 20% substrate depletion, the concentration of CYP enzyme, Fdx and FdR used was 1 µM, 10 µM and 0.05 U/mL, respectively. The 11-hydroxylauric acid metabolite was quantified based on a calibration curve prepared in matrix and ranging from 0.1 µM to 10 µM. The mean metabolite formation rate values obtained from triplicate determinations were fit to the Michaelis-Menten (hyperbolic) and Hill (sigmoidal) equations using GraphPad Prism software (version 9.0.0). Comparison of the best fit was based on the second order Aikake Information Criterion (AICc) analysis (Table 2).

### Lauric Acid Metabolite Analysis by GC-MS

Lauric acid metabolites generated in incubations of 100 µM lauric acid with CYP168A1 (5 µM) and the spinach redox partners Fdx (5 µM) and FdR (0.13 U/mL) were extracted twice with dichloromethane and dried at room temperature under nitrogen flow. Samples, resuspended in acetonitrile, were derivatized by addition of 50 µL *N*-methyl-*N*-trimethylsilyl-trifluoroacetamide (containing trimethylchlorosilane at 1% v/v) followed by 20 min incubation at 70°C. Derivatized samples were transferred to Teflon capped vials for GC-MS analysis on an Agilent Technologies 5977/7890 gas chromatograph using an Agilent HP-5MS column (30 m × 0.25 mm inside diameter × 0.25 µm). Separation of trimethylsilyl-derivatized lauric acid and hydroxyl metabolites was achieved by temperature gradient as following: 70°C for 1 min, increased 25°C/min up to 170°C, then increased 5°C/min up to 200°C and increased 20°C/min up to 280°C, held for 5 min at 280°C. GC parameters were as follows: inlet temperature of 250°C, splitless constant flow mode at 7.57 mL/min, and MS transfer line at 230°C. Lauric acid and its metabolites were ionized using electron impact ionization and detected by a single quadrupole in scan mode from 30-500 mass units. MS parameters were as follows: source temperature of 230°C, quadrupole temperature of 150°C. The GC–MS system was controlled by an Agilent MassHunter Workstation. Data was analyzed using Agilent MassHunter Quantitative Analysis. Identification of the trimethylsilyl-derivatized 11-hydroxylauric acid was confirmed using an authentic standard and verified using the NIST17 GCMS mass spectral database (115).

### LC-MS Method for Lauric Acid Hydroxylation

The lauric acid incubation samples with the recombinant CYP168A1 enzyme were analyzed by LC-MS with a Waters Acquity Ultra-Performance Liquid Chromatography (UPLC) system interfaced by electrospray ionization with a Waters Xevo TQ-S micro tandem quadrupole mass spectrometer (Waters Corp., Milford, MA) in negative ionization mode and with multiple reaction monitoring (MRM) scan type. Due to limited fragmentation of lauric acid, its metabolites and the internal standard, a parent-to-parent mass transition strategy was employed. The following mass transitions, collision energies (CEs), and cone voltages (CVs) were used to detect the respective analytes: 199.1>199.1, CE = 10 V, CV = 40 V for lauric acid, 215.1>215.1, CE = 10 V, CV = 40 V for the hydroxylauric acid metabolites and 235.2>235.2, CE = 8 V, CV = 20 V for the internal standard 12-hydroxylauric-d_20_ acid. The following source conditions were applied: 1 kV for the capillary voltage, 150°C for the source temperature, 500°C for the desolvation temperature and 900 L/h for the desolvation gas flow. Lauric acid and its hydroxylated metabolites were separated on a Waters BEH C18 column (1.7 µm, 2.1 x 50 mm) by flowing 2 mM ammonium acetate in water and in methanol at 0.4 mL/min. The following gradient was used: 10% organic (methanol) held for 0.5 min, increased first to 45% over 0.5 min, then increased to 53% over 2 min, and finally increased to 98% over 0.2 min and held at 98% over 1.8 min. To limit soiling of the source, a divert directing the LC flow to waste was set at 3.5 min before elution of lauric acid. The MS peaks were integrated using QuanLynx software (version 4.1, Waters Corp., Milford, MA), and the analyte/internal standard peak area ratios were used for relative quantification. For determination of the hydroxy metabolite concentration, the regression fit was based on the analyte/internal standard peak area ratios calculated from the calibration standards, and the analyte concentration in the incubations was back-calculated using the weighted (1/x) linear least-squares regression.

### Recombinant CYP168A1 Arachidonic Acid in Vitro Metabolic Assays

The same CYP:Fdx:FdR ratio used for lauric acid kinetic experiments was initially used for assessing linearity in arachidonic acid metabolism over 60 min. However, under these conditions, metabolite linearity couldn’t be established and decrease in CYP168A1, Fdx and FdR concentrations was necessary. The CYP:Fdx:FdR ratio of 0.5 µM:5 µM:0.025 U/mL allowed to achieve product formation linearity up to 20 min and to stay under steady state conditions. Incubations of arachidonic acid with CYP168A1 and the spinach redox partners were done with slight modifications of McDonald et al. (116). Briefly, arachidonic acid (sodium salt) prepared in methanol to yield final concentrations between 2.5 to 200 µM in the reactions was incubated with CYP168A1 and the spinach redox partners in 100 mM potassium phosphate (pH 7.4), 3 mM MgCl_2_ and 1 mM sodium pyruvate. After an equilibration at 37°C for 3 min, the reactions, prepared in triplicate, were initiated by the addition of a NADPH-regenerating system mix consisting of NADP^+^ (1 mM), D-glucose-6-phosphate (10 mM) and glucose-6-phosphate dehydrogenase (2 IU/mL). After 20 min at 37°C and under agitation, the enzymatic reactions, done in dim conditions, were stopped by the addition of ice-cold methanol (200 µL) containing 600 ng/mL 20-HETE-d_6_ internal standard and 0.02% 2,6-di-tert-butyl-4-methylphenol. Incubations without the NADPH-regenerating system mix served as negative controls. For the arachidonic acid incubations with ketoconazole, a ketoconazole stock solution was made in methanol and was added in the CYP168A1 reactions prepared in triplicate to yield a final concentration of 10 µM ketoconazole. Methanol solvent control incubations were done in parallel. Precipitated proteins were collected by centrifugation of the stopped reaction samples at 2,500 × *g* and 4 °C for 20 min. Supernatants were transferred to HPLC vials, and aliquots of 5 µL were analyzed by LC-MS. The 18-HETE and 19-HETE metabolites were quantified based on calibration curves prepared in matrix and ranging from 0.05 µM to 10 µM and 0.1 µM to 10 µM, respectively. The 19(S)-HETE standard was used to prepare the calibration curve, since no racemic mixture of 19-HETE was commercially available.

### LC-MS Method for Arachidonic Acid Hydroxylation

The arachidonic acid incubation samples with the recombinant CYP168A1 enzyme were analyzed by LC-MS with a Waters Acquity Ultra-Performance Liquid Chromatography (UPLC) system interfaced by electrospray ionization with a Waters Xevo TQ-S micro tandem quadrupole mass spectrometer (Waters Corp., Milford, MA) in negative ionization mode and with multiple reaction monitoring (MRM) scan type. The following source conditions were applied: 1 kV for the capillary voltage, 150°C for the source temperature, 500°C for the desolvation temperature and 900 L/h for the desolvation gas flow. The following mass transitions, collision energies (CEs), and cone voltages (CVs) were used to detect the respective analytes: 303.1>259.1, CE = 12 V, CV = 20 V for arachidonic acid, 319.0>261.3, CE = 18 V, CV = 20 V for the 18-HETE, 319.0>231.3, CE = 15 V, CV = 20 V for the 19-HETE, 319.0>289.3, CE = 15 V, CV = 20 V for the 20-HETE and 325.1>281.2, CE = 15 V, CV = 20 V for the internal standard 20-HETE-d_6_. Arachidonic acid and its hydroxylated metabolites were separated on a Waters BEH C18 column (1.7 µm, 2.1 x 100 mm) by flowing 2 mM ammonium acetate in water and in 4:1 acetonitrile:methanol at 0.3 mL/min. The following gradient was used: 55% organic (4:1 acetonitrile:methanol) held for 3.5 min, increased to 98% over 0.5 min, and held at 98% over 2 min. The MS peaks were integrated using QuanLynx software (version 4.1, Waters Corp., Milford, MA), and the analyte/internal standard peak area ratios were used for relative quantification. For determination of the hydroxy metabolite concentration, the regression fit was based on the analyte/internal standard peak area ratios calculated from the calibration standards, and the analyte concentration in the incubations was back-calculated using the weighted (1/x) linear least-squares regression. For acquisition of the HETE metabolite MSMS spectra, daughter scans of *m/z* 319 were acquired in centroid mode between *m/z* of 50 to 350 using the above source conditions and a collision energy of 18 V.

### CYP168A1 Homology Model Construction

A BlastP search of the CYP168A1 sequence (NP_251165) revealed that the top scoring hit is a protein known as CalO2, a putative CYP enzyme from *Micromonospora echinospora*, generating a MAX score of 145. However, this organism is not closely related to *P. aeruginosa*. Therefore, the more closely related second top scoring hit CYP P450 Vdh, from *P. autotrophic*, was used as a template for building the homology model of CYP168A1. Both CYP168A1 and CYP P450 Vdh were used as target and template, respectively. Due to the lack of similarity of the first 21 amino acids to any known sequence in the PDB or NCBI RefSeq database, these residues were omitted for the purposes of homology model construction. This resulted in greater than 90% sequence homology between CYP168A1 and CYP P450 Vdh. Models were generated using UCSF MODELLER (50) ran locally via the UCSF Chimera GUI interface (117). MODELLER parameters included an output of 5 independent models, non-water HETATM residues from the template (heme), and hydrogen atoms. The final model chosen produced a zDOPE score, an atomic distance-dependent statistical score where negative values indicate better models, of -0.2. Additionally, the model had a GA341 score of 1. The GA341 score is a model score derived from statistical potentials where a value >0.7 generally indicates a reliable model, i.e., >95% probability of having the correct fold. Following initial model generation, unstructured loops were refined using the loops refinement MODELLER plugin in Chimera. After loop refinement, the zDOPE score was reduced to -1.08 and the GA341 score remained unchanged. This final refined model was used for all subsequent ligand docking studies.

### Docking of Substrates and Ketoconazole to CYP168A1

In order to understand how substrate and inhibitor ligands structurally interact with CYP168A1, an in silico docking study was undertaken using AutoDock Vina (53) and the CYP168A1 homology model as the receptor template. The protein was prepared for docking using MGLTools, AutoDock Tools V. 1.5.7 (The Scripps Research Institute, USA) by adding polar hydrogens and assigning partial charges. Coordinates for the docking grid search space were established by defining the enzyme active site, with the final parameters being: grid box center; x-center = 9.285, y-center = 21.861, z-center = 19.061, and the total number of grid points in each dimension being; x-dimension = 24.664, y-dimension = 29.966, and z-dimension = 29.793. The ligands selected for docking were ketoconazole, a Type II inhibitor, and the fatty acid substrates, lauric acid and arachidonic acid. Each ligand file was downloaded from the Protein Data Bank (PDB: https://www.rcsb.org/) and parameterized for docking in the following manner: 1) addition of polar hydrogens, 2) assessment (and assignment, when necessary) of rotatable bonds, and 3) assignment of partial charges. Both receptor (protein) and ligand files were saved in the PDBQT format. A configuration file docking script was prepared in simple text format with the energy range set to 4 and the exhaustiveness search parameter set to 8. AutoDock Vina was invoked using the configuration file and PDBQT.out and log.out files. Output files were qualitatively and quantitatively analyzed by using the VewDock function of UCSF Chimera, and ranked by binding energy (ΔG). The most energetically favorable binding mode for each ligand was reported as 0 rmsd. All figures were generated using UCSF Chimera (117).

## Supporting information

CYP168A1-homology-model

## DATA AVAILABILITY

All data is made publicly available through the JBC repository or may be obtained by contacting the corresponding author directly.

## SUPPORTING INFORMATION

The homology model for CYP168A1 used in this study is made available as Supporting Information.

## ACKNOWLEDGEMENTS

We would like to gratefully acknowledge Michael Armstrong of the CU Skaggs School of Pharmacy and Pharmaceutical Sciences Mass Spectroscopy facility for assistance provided in confirming the CYP168A1 lauric acid metabolites.

## AUTHOR CONTRIBUTIONS

B.C.T., H.M.W., S.E.K., and J.N.L. participated in research design and writing and editing the manuscript. B.C.T., H.M.W., S.E.K. participated in conducting experiments, data collection, and B.C.T., H.M.W., S.E.K., and J.N.L. conducted data analysis. J.N.L directed all aspects of the research conducted.

## FUNDING AND ADDITIONAL INFORMATION

The research described in this manuscript was generously funded through a University of Colorado, Skaggs School of Pharmacy and Pharmaceutical Sciences Faculty Start-up Package and the 2021-22 Skaggs Scholar Award.

## ABBREVIATIONS AND NOMENCLATURE

